# Voluntary oxycodone self-administration produces analgesic tolerance and sex-dependent hyperalgesia across genetically diverse rats

**DOI:** 10.64898/2026.08.20.745912

**Authors:** Tolulope J. Ajanaku, Eamonn P. Duffy, Jonathon O. Ward, Luanne H. Hale, Caleb I. Hodges, Laura M. Saba, Marissa A. Ehringer, Ryan K. Bachtell

## Abstract

Long-term opioid therapy is limited by analgesic tolerance and opioid-induced hyperalgesia, but the roles of genetic background, sex, and drug exposure remain unclear. We used 20 inbred strains from the Hybrid Rat Diversity Panel to examine thermal sensitivity, oxycodone analgesia, tolerance, and hyperalgesia-like changes following voluntary intravenous oxycodone or saline self-administration. Rats underwent tail-immersion testing before self-administration (Pre-SA) and after self-administration (Post-SA). Oxycodone analgesia was assessed using the percent maximum possible effect time course and the corresponding area under the curve. Pre-SA thermal sensitivity differed across strains and between sexes, and Pre-SA oxycodone analgesia also differed across strains. Oxycodone self-administration produced a sex-dependent increase in thermal sensitivity that was most evident in males. During Post-SA testing, oxycodone self-administering rats showed reduced analgesic responsiveness compared with saline controls, and the magnitude of this difference varied across strains. Within-strain Pre-SA-to-Post-SA comparisons identified tolerance-like reductions in several strains. Across strains and sexes, oxycodone self-administering rats showed a greater Pre-SA-to-Post-SA reduction in analgesic responsiveness than saline controls, consistent with analgesic tolerance. Total oxycodone intake was not associated with tolerance at either the strain-mean or individual-animal level. Heritability estimates were higher for thermal sensitivity and analgesia (H² ≈ 0.28–0.40) than for changes in thermal sensitivity and tolerance (H² ≈ 0.18–0.27). These findings demonstrate strain variation in thermal sensitivity and oxycodone analgesia, sex-dependent hyperalgesia-like effects, and reduced analgesic responsiveness following voluntary oxycodone intake.

## Introduction

Prescription opioids such as oxycodone are widely used to treat moderate to severe pain, but their long-term use is limited by the development of analgesic tolerance, opioid-induced hyperalgesia (OIH) and opioid use disorder (OUD) [1–5]. These complications mean that patients often require increasing doses to achieve the same level of pain relief, which increases the risk of side effects, overdose, and addiction. Both humans and animal models show marked individual differences in baseline pain sensitivity and initial opioid analgesic responsiveness [6–9]. In genetically diverse rodent populations, this variability also extends to voluntary oxycodone intake and susceptibility to analgesic tolerance and OIH [10–12]. However, the contributions of genetic background and other biological factors to this variability are still not well understood [13].

A major ongoing question in opioid research is how repeated opioid exposure produces analgesic tolerance and OIH [3,10]. These adaptations are important because they reduce analgesic efficacy and drive dose escalation. They are also thought to contribute to the transition from controlled opioid use to compulsive drug seeking [14]. While human genome-wide association studies can identify genetic variants associated with opioid-related clinical outcomes, they are generally limited in their ability to assess detailed experimentally controlled phenotypes such as analgesic tolerance and hyperalgesia [15–17]. The Hybrid Rat Diversity Panel helps address this limitation by allowing multiple pain-related and opioid self-administration phenotypes to be measured in the same rats under controlled experimental conditions [7,11]. Consequently, the present study used 20 inbred rat strains from the HRDP to investigate individual and strain differences in voluntary oxycodone intake and in the development of tolerance and hyperalgesia over long-term exposure under controlled conditions.

Studies using genetically defined rodent populations have established that individual differences in opioid analgesia [7,9,12,18,19], analgesic tolerance [12,20], and OIH [10] are influenced by genetic background. For example, Kest et al. observed pronounced strain differences in morphine analgesic tolerance across 11 inbred mouse strains, with some strains developing robust tolerance and others showing little or no tolerance [20]. Similarly, Yang et al. reported substantial variation in morphine analgesic responsiveness across 15 inbred Genetic Diversity mouse strains, further demonstrating that genetic background contributes to individual differences in opioid response [18]. More recently, work using the HRDP showed that both genetic background and sex significantly influence baseline somatosensory sensitivity and oxycodone analgesia [7]. Together, these studies underscore the critical role of genetic background in shaping opioid-related phenotypes and provide a foundation for using genetically defined rodent panels to dissect the mechanisms underlying individual differences in opioid tolerance and hyperalgesia.

The overall aim of the present study was to determine whether baseline thermal sensitivity, acute oxycodone analgesia, analgesic tolerance, and OIH-like changes differed by genetic background and sex in 20 HRDP inbred rat strains following voluntary oxycodone or saline self-administration (SA). Using a voluntary intravenous SA paradigm combined with Pre-SA and Post-SA tail-immersion testing, we also examined whether the magnitude of analgesic tolerance was related to total oxycodone intake. This approach identified substantial strain variation in thermal sensitivity and oxycodone analgesia, sex-dependent hyperalgesia following oxycodone SA, and an overall reduction in analgesic responsiveness consistent with tolerance.

## Methods

### Subjects

Adult (PND 60+) male and female rats (n=615) from 20 HRDP strains were used for the current study (Table 1). Two strains (F344/NCrl and WKY/NCrl) were obtained from Charles River Laboratories and LEW/SSNHsd was obtained from ENVIGO. The remaining 17 strains were obtained from the Medical College of Wisconsin (provided by Dr. Melinda Dwinell, R24OD024617). Upon arrival, rats were habituated for at least one week. Rats were individually housed at 22°C and 40% humidity under a 12-h light/12-h dark cycle. Rats were provided with standard rat chow (Teklad 2918, Envigo, Indianapolis, IN) and water *ad libitum*. The ACI/EurMcwi strain was included in all batches obtained from the Medical College of Wisconsin as a common reference strain across batches. This strain was chosen due to preliminary studies identifying that the ACI/EurMcwi strain displays robust behavioral responses in all outcome measures. All procedures involving animals were conducted in accordance with institutional and federal guidelines and approved by the Institutional Animal Care and Use Committee at the University of Colorado Boulder (Protocol 2742), an Association for Assessment and Accreditation of Laboratory Animal Care International-accredited institution.

**Table 1.** Sample sizes of animals used per strain, sex, and self-administration group. The number of male and female rats from each HRDP strain assigned to the oxycodone (OXY) or saline (SAL) self-administration groups. Strains, Research Resource Identifier (RRID) accession numbers, and origin facilities are indicated. Totals are shown for each sex, strain, self-administration group, and for the overall cohort (*n* = 615; females = 304, males = 311). CRL, Charles River Laboratories; MCW, Medical College of Wisconsin; TOT, total number of rats.

| STRAIN | RRID | VENDOR | FEMALE |  |  | MALE |  |  | GRAND |
| --- | --- | --- | --- | --- | --- | --- | --- | --- | --- |
|  |  |  | OXY | SAL | TOT | OXY | SAL | TOT |  |
| ACI/EurMcwi | RRRC_00284 | MCW | 26 | 15 | 41 | 27 | 15 | 42 | 83 |
| BN-Lx/Cub | RGD_61117 | MCW | 5 | 5 | 10 | 7 | 5 | 12 | 22 |
| BXH2/Cub | RGD_2307121 | MCW | 12 | 9 | 21 | 6 | 8 | 14 | 35 |
| BXH6/Cub | RGD_2307136 | MCW | 10 | 6 | 16 | 12 | 7 | 19 | 35 |
| BXH12/Cub | RGD_2307139 | MCW | 5 | 3 | 8 | 5 | 3 | 8 | 16 |
| F344/NCrl | RGD_737926 | CRL | 9 | 8 | 17 | 9 | 9 | 18 | 35 |
| F344/Stm | RGD_1302686 | MCW | 7 | 6 | 13 | 12 | 10 | 22 | 35 |
| HXB1/Ipcv | RGD_2307094 | MCW | 8 | 5 | 13 | 8 | 4 | 12 | 25 |
| HXB2/IpcvMcwi | RGD_14973552 | MCW | 7 | 5 | 12 | 9 | 5 | 14 | 26 |
| HXB10/IpcvMcwi | RGD_38549351 | MCW | 7 | 6 | 13 | 9 | 7 | 16 | 29 |
| HXB17/Ipcv | RGD_2307085 | MCW | 6 | 3 | 9 | 8 | 6 | 14 | 23 |
| HXB18/Ipcv | RGD_2307082 | MCW | 4 | 3 | 7 | 3 | 2 | 5 | 12 |
| HXB23/Ipcv | RGD_2307093 | MCW | 14 | 11 | 25 | 11 | 10 | 21 | 46 |
| HXB31/IpcvMcwi | RGD_14973551 | MCW | 8 | 5 | 13 | 7 | 5 | 12 | 25 |
| LE/Stm | RGD_629485 | MCW | 8 | 7 | 15 | 5 | 7 | 12 | 27 |
| LEW/Crl | RGD_737932 | CRL | 7 | 5 | 12 | 7 | 5 | 12 | 24 |
| LEW/SSNHsd | RGD_737922 | ENVIGO | 7 | 5 | 12 | 6 | 6 | 12 | 24 |
| M520/N | RRRC_00168 | MCW | 10 | 7 | 17 | 9 | 6 | 15 | 32 |
| SHR/OlaIpcv | RGD_631848 | MCW | 6 | 6 | 12 | 7 | 6 | 13 | 25 |
| WKY/NCrl | RGD_1358112 | CRL | 10 | 8 | 18 | 9 | 9 | 18 | 36 |
| **Total** |  |  | **176** | **128** | **304** | **176** | **135** | **311** | **615** |

Values are numbers of rats. OXY, oxycodone self-administration group; SAL, saline self-administration group; TOT, total; CRL, Charles River Laboratories; MCW, Medical College of Wisconsin; RRID, Research Resource Identifier.

### Study design

The study consisted of four sequential phases: Pre-SA behavioral testing, catheter surgery, voluntary self-administration (SA), and Post-SA behavioral testing. During the Pre-SA phase, rats underwent tail-immersion testing to assess thermal sensitivity and acute oxycodone analgesia, and von Frey testing to assess mechanical sensitivity. The Pre-SA von Frey data have been reported previously [7]. After Pre-SA testing, rats underwent catheter surgery before beginning SA. Rats were then assigned to either the Oxy-SA group or the Sal-SA group and completed a 20-day self-administration protocol. After completion of SA, rats underwent Post-SA behavioral testing under the same general conditions used during the Pre-SA assessment. The present manuscript focuses on tail-immersion outcomes, including thermal sensitivity, oxycodone analgesia, and analgesic tolerance, with Sal-SA rats serving as time-and procedure-matched controls.

### Tail-immersion test

The tail-immersion test was conducted both before (Pre-SA) and after (Post-SA) the SA period (raw rat-level tail-immersion data in S1 Data). Post-SA tail-immersion testing was conducted 2 days after the final SA session. This assay involved immersing 5–10 cm of the distal portion of the rat’s tail into a hot water bath maintained at 49 ± 1°C and recording the latency (s) for the rat to generate a tail-flick or tail-withdrawal response. The maximum cutoff latency was set at 15 s. These parameters were selected based on preliminary studies indicating that longer cutoff times and temperatures above 50°C produced tissue damage in some inbred strains. Tail-withdrawal latencies were recorded at 0, 15, 30, 45, 60, 90, and 120 min. The 0-min timepoint was used to evaluate each rat’s initial thermal sensitivity. Immediately following the 0-min measurement, rats received acute oxycodone (1 mg/kg, i.p.). Repeated measurements at subsequent timepoints were used to assess the development, magnitude, and duration of oxycodone-induced analgesia. Analgesia typically increased shortly after oxycodone exposure, peaked during the early post-injection period (15–30 min), and gradually returned toward baseline by 120 min. Although latencies were collected through 120 min, analyses focused on the 0–60 min %MPE time course and its corresponding AUC.

### Surgery

Chronic indwelling intra-jugular catheters were implanted under aseptic conditions as described previously [11]. Rats were anesthetized with isoflurane (2–4%) and received carprofen (5 mg/kg, s.c.) and enrofloxacin (5 mg/kg, s.c.) immediately prior to surgery. Postoperative analgesia consisted of carprofen (5 mg/kg, s.c.) administered daily for two days following surgery. Rats were allowed 7 days to recover before beginning self-administration procedures.

### Self-administration procedures

Prior to SA, rats within each strain were assigned to saline or oxycodone SA groups using a pseudorandomized procedure based on Pre-SA tail-immersion testing to ensure balanced initial thermal sensitivity and oxycodone analgesic responsiveness across future SA groups. This balance was confirmed statistically, with no significant differences between rats assigned to the oxycodone SA (Oxy-SA) and saline SA (Sal-SA) groups in either baseline tail-withdrawal latency or Pre-SA oxycodone analgesia (both p > 0.05). SA sessions took place in operant chambers (Med Associates, St. Albans, VT) equipped with two retractable levers along with cue lights above each lever and a sound-attenuating fan. All sessions occurred at dark-cycle onset (12-h light:12-h dark). Acquisition consisted of ten 2-h short-access sessions on an FR1 schedule, during which a response on the active lever delivered a 5-s saline or oxycodone infusion (0.15 mg/kg/infusion; Oxycodone HCl, B&B pharmaceuticals, Englewood, CO) paired with a 20-s cue light (7.5-W white) and followed by a 20-s timeout; inactive-lever responses were recorded but produced no programmed consequence. Following acquisition and a brief intervening progressive-ratio test, rats entered the escalation phase — ten 12-h long-access sessions under the same FR1:TO20 contingencies. SA here served solely as the exposure manipulation for assessing analgesic tolerance; full SA behavioral outcomes are reported elsewhere [11].

### Behavioral Phenotypes

Table 2 provides an overview of the definitions, calculations, and interpretation of behavioral phenotypes used to assess various phenotypes.

**Table 2.** Definitions, calculations, and interpretation of behavioral phenotypes assessed before and after self-administration.

| Phenotype | Measurement or calculation | Interpretation |
| --- | --- | --- |
| **Pre-SA thermal sensitivity** | Pre-SA tail withdrawal latency at the 0-min timepoint | Shorter latency indicates greater thermal sensitivity |
| **Post-SA thermal sensitivity** | Post-SA tail withdrawal latency at the 0-min timepoint | Shorter latency indicates greater thermal sensitivity |
| **Opioid-induced hyperalgesia** | Post-SA 0-min latency – Pre-SA 0-min latency | Negative values indicate increased thermal sensitivity after self-administration |
| **Pre-SA oxycodone analgesia** | **Pre-SA %MPE time course:** 15, 30, 45, and 60 min following the Pre-SA oxycodone challenge | Represents the time-dependent analgesic response to Pre-SA oxycodone |
|  | **Pre-SA %MPE AUC:** Area under the Pre-SA %MPE–time curve following Pre-SA oxycodone challenge | Higher values indicate greater magnitude and duration of Pre-SA oxycodone analgesia |
| **Post-SA oxycodone analgesia** | **Post-SA %MPE time course:** 15, 30, 45, and 60 min following the Post-SA oxycodone challenge | Represents the time-dependent analgesic response to Post-SA oxycodone |
|  | **Post-SA %MPE AUC:** Area under the Post-SA %MPE–time curve following Post-SA oxycodone challenge | Higher values indicate greater magnitude and duration of Post-SA oxycodone analgesia |
| **Tolerance** | **Tolerance AUC Difference:** Post-SA %MPE AUC – Pre-SA %MPE AUC | Negative values indicate reduced analgesic responsiveness after self-administration |
| **Self-administration intake** | **Total Oxycodone Intake:** Oxycodone infusions received during the 20-day self-administration phase multiplied times unit dose (0.15 mg/kg) | Defines amount of chronic oxycodone exposure |

AUC, area under the curve; %MPE, percent maximum possible effect; Pre-SA, pre-self-administration; Post-SA, post-self-administration

#### Thermal sensitivity

Thermal sensitivity was assessed using raw tail-withdrawal latencies measured at the 0-min timepoint during the Pre-SA and Post-SA tests. Change in thermal sensitivity was calculated as Post-SA minus Pre-SA 0-min tail-withdrawal latency, with negative values indicating increased thermal sensitivity. The Pre-SA baseline analysis included rats with a valid Pre-SA 0-min measurement, whereas the change-score analysis included only rats with valid 0-min measurements at both Pre-SA and Post-SA.

#### Oxycodone analgesia

Pre-SA and Post-SA oxycodone analgesia was evaluated using two complementary measures: the percent maximum possible effect (%MPE) time course and the area under the %MPE–time curve from 0 to 60 min. Tail-withdrawal latencies were recorded following acute oxycodone administration before and after the self-administration period. Latencies were converted to %MPE to express the oxycodone-induced response relative to each rat’s baseline latency and the 15-s assay cutoff, thereby reducing the influence of baseline differences in thermal sensitivity and facilitating comparisons of analgesic responsiveness across animals and strains. %MPE was calculated as:

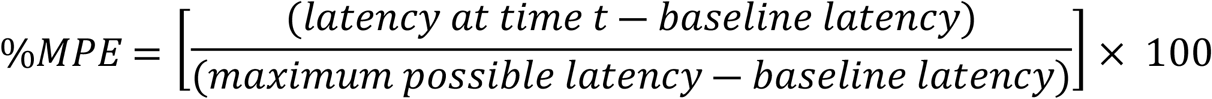

where baseline latency was the 0-min latency for that testing session and the maximum possible latency was 15 s. Because each session was normalized to its own baseline, the 0-min value equaled 0% MPE by definition and was retained as the zero anchor in the time-course analyses, figures, and 0–60-min AUC calculations. %MPE values were bounded to 0–100%, with values below 0 set to 0 and values above 100 set to 100, to keep analgesic responses within a biologically interpretable range and avoid outlier influence.

A 0-min baseline latency equal to the 15-s assay cutoff occurred in three Pre-SA sessions (ACI/EurMcwi strain, n=2 and SHR/OlaIpcv strain, n=1) and three Post-SA sessions (ACI/EurMcwi strain, n=2 and HXB17/Ipcv strain, n=1), resulting in a zero denominator for the %MPE calculation at those sessions. Undefined %MPE values were not included.

Pre-SA and Post-SA %MPE time courses were analyzed directly. Corresponding 0–60-min AUCs were calculated using trapezoidal integration of %MPE values at 0, 15, 30, 45, and 60 min. Complete data at all five timepoints were required, and rats with missing values at any timepoint were excluded from the corresponding AUC analysis.

#### Analgesic tolerance

Analgesic tolerance was evaluated using complementary between-group and within-subject approaches. The between-group approach assessed whether rats in the Oxy-SA group showed reduced Post-SA oxycodone analgesic responsiveness compared with rats in the Sal-SA group. Between-group tolerance was evaluated using the Post-SA %MPE time-course profiles for the Oxy-SA and Sal-SA groups. The within-subject approach assessed whether oxycodone analgesic responsiveness declined from Pre-SA to Post-SA within the same animals. Within-subject tolerance was evaluated using both the Pre-SA and Post-SA %MPE time-course profiles and the Tolerance AUC Difference. The Tolerance AUC Difference was calculated as follows:

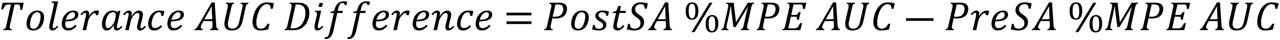

Negative Tolerance AUC Difference values indicated reduced analgesic responsiveness from Pre-SA to Post-SA in both groups. A greater negative change in the Oxy-SA group than in the Sal-SA group was interpreted as evidence of analgesic tolerance associated with oxycodone self-administration; values in Sal-SA rats controlled for changes resulting from procedural factors.

#### Total oxycodone intake

Total oxycodone intake was calculated for each oxycodone self-administering rat by summing all oxycodone-reinforced infusions across self-administration sessions and multiplying by the unit dose of 0.15 mg/kg per infusion. Total intake was expressed in mg/kg and was used to examine whether the amount of oxycodone consumed was associated with the magnitude of analgesic tolerance.

### Statistical analyses

Data are reported as mean ± standard error of the mean (SEM) unless otherwise stated. Behavioral data (raw tail withdrawal latencies, percent maximum possible effect [%MPE], and area under the curve [AUC] measures) were processed and analyzed in R (version 4.5.2) using the packages dplyr, readxl, readr, purrr, tidyr, stringr, ggplot2, viridis, lme4, lmerTest, emmeans, and pracma. Statistical significance was set at p < 0.05. Post hoc comparisons were conducted using estimated marginal means from the emmeans package, with Bonferroni adjustment for multiple comparisons where appropriate.

#### Thermal sensitivity analyses

Thermal sensitivity was assessed using raw tail-withdrawal latency at the 0-min timepoint. Pre-SA 0-min tail-withdrawal latency was analyzed using a two-way ANOVA with strain, sex, and their interaction as factors. Change in thermal sensitivity was calculated as Post-SA 0-min latency minus Pre-SA 0-min latency, with negative values indicating increased thermal sensitivity after self-administration. Change in thermal sensitivity was analyzed using a three-way ANOVA with strain, sex, self-administration group, and their interactions as factors. Post hoc comparisons were conducted using estimated marginal means where appropriate.

#### Oxycodone analgesia analyses

%MPE time course data were analyzed using repeated-measures ANOVA implemented as linear mixed-effects models. Timepoint was treated as a categorical within-subject factor, and strain, sex, and self-administration group were treated as between-subject factors where applicable. Models were fit using the lmer function, with rat ID included as a random intercept to account for repeated measurements within animals. Fixed-effect tests were obtained using lmerTest.

For Pre-SA oxycodone analgesia, the %MPE time course was analyzed using a linear mixed-effects model with strain, sex, timepoint, and all corresponding interactions included as fixed effects. Rat ID was included as a random intercept. Pre-SA %MPE AUC was analyzed separately using a two-way ANOVA with strain, sex, and their interaction as factors.

For Post-SA oxycodone analgesia, the %MPE time course was first analyzed using a full linear mixed-effects model with strain, self-administration group, timepoint, sex, and their interactions included as fixed effects, with rat ID included as a random intercept. Sex and sex-related interaction terms were not significant in the full model. A likelihood-ratio test comparing the full model with a reduced model excluding sex-related terms indicated that sex-related terms did not improve model fit. Therefore, the final Post-SA time-course model included strain, self-administration group, timepoint, and their interactions as fixed effects, with rat ID included as a random intercept.

Post-SA %MPE AUC also was first analyzed using a full factorial ANOVA with strain, sex, self-administration group, and their interactions as factors. Sex and sex-related interaction terms were not significant in the full model. A model comparison indicated that sex-related terms did not improve model fit. Therefore, the final Post-SA %MPE AUC model included strain, self-administration group, and their interaction as factors.

#### Analgesic tolerance analyses

Analgesic tolerance was assessed using complementary between-group and within-subject approaches. For the between-group analysis, the dependent measure was Post-SA %MPE across the 0, 15, 30, 45, and 60 min timepoints. Post-SA %MPE was analyzed using a linear mixed-effects model with strain, self-administration group, timepoint, and their interactions as fixed effects and rat ID as a random intercept. This analysis tested whether rats in the Oxy-SA group showed reduced Post-SA analgesic responsiveness compared with rats in the Sal-SA group and whether this difference varied across strains and timepoints.

Within-subject changes in analgesic responsiveness were evaluated in two complementary ways. First, Pre-SA and Post-SA %MPE time courses were compared within the Oxy-SA or Sal-SA group using separate linear mixed-effects model with phase, strain, timepoint, and their interactions as fixed effects and rat ID as a random intercept. Phase was defined as Pre-SA versus Post-SA. Pre-SA and Post-SA estimated marginal means were compared at 15, 30, 45, and 60 min within each strain, with Bonferroni correction applied across the four post-injection timepoints separately within each strain. The 0-min timepoint was excluded from these comparisons because %MPE was zero by definition.

Second, a Tolerance AUC Difference score was calculated for each rat as Post-SA %MPE AUC minus Pre-SA %MPE AUC. Negative values indicated lower Post-SA analgesic responsiveness relative to Pre-SA, consistent with greater tolerance-like reduction. These within-rat difference scores were analyzed using factorial ANOVA with strain, sex, self-administration group, and their interactions included as fixed effects.

### Correlational analyses

To assess the relationship between oxycodone exposure and tolerance, Pearson correlation coefficients were calculated between Tolerance AUC difference scores and total oxycodone intake (mg/kg). Correlations were examined at both the individual-animal level, and the level of strain means. As a post hoc sensitivity analysis, the association between strain-mean total oxycodone intake and the strain-mean 15-min tolerance difference, calculated as Post-SA %MPE minus Pre-SA %MPE, was also examined using Pearson correlation.

#### Heritability

Broad-sense heritability (H²) for each behavioral phenotype was estimated using the coefficient of determination (R²) from a two-way ANOVA, following the approach previously used in the HRDP [7]. For the present analyses, the model included strain, sex, and the strain × sex interaction. The proportion of phenotypic variance explained by the full model was used as the H² estimate for each phenotype.

## Results

### Thermal sensitivity

A two-way ANOVA was used to examine the effects of strain, sex, and their interaction on Pre-SA baseline thermal sensitivity (S1 Fig). At the Pre-SA 0-min timepoint, tail-withdrawal latency differed significantly across strains (F₁₉,₅₇₅ = 11.72, p < 0.001) and between sexes (F₁,₅₇₅ = 9.65, p = 0.002). Estimated marginal means showed that males had longer baseline tail-withdrawal latencies than females (p = 0.002), indicating greater baseline thermal sensitivity in females. The strain × sex interaction was not significant (p = 0.090).

To quantify oxycodone-induced hyperalgesia-like changes, we analyzed the change in 0-min tail-withdrawal latency from Pre-SA to Post-SA, with negative values indicating increased thermal sensitivity (Fig 1). There was a significant main effect of strain (F₁₉,₄₂₇ = 2.53, p < 0.001), indicating that the magnitude of change in thermal sensitivity differed across strains. The strain × sex, strain × SA group, and strain × sex × SA group interactions were not significant. However, there was a significant sex × SA group interaction (F₁,₄₂₇ = 7.43, p = 0.0067), indicating that the effect of SA group on change in thermal sensitivity differed by sex. Post hoc comparisons showed no difference between the Sal-SA and Oxy-SA groups in females (t₄₂₇ = 0.28, p = 0.78). In males, the Oxy-SA group showed a greater reduction in 0-min tail-withdrawal latency than the Sal-SA group (t₄₂₇ = 4.02, p < 0.001). These results indicate that change in thermal sensitivity varied across strains and that oxycodone SA produced a sex-dependent increase in thermal sensitivity, with the effect most evident in males.

**Fig 1.**
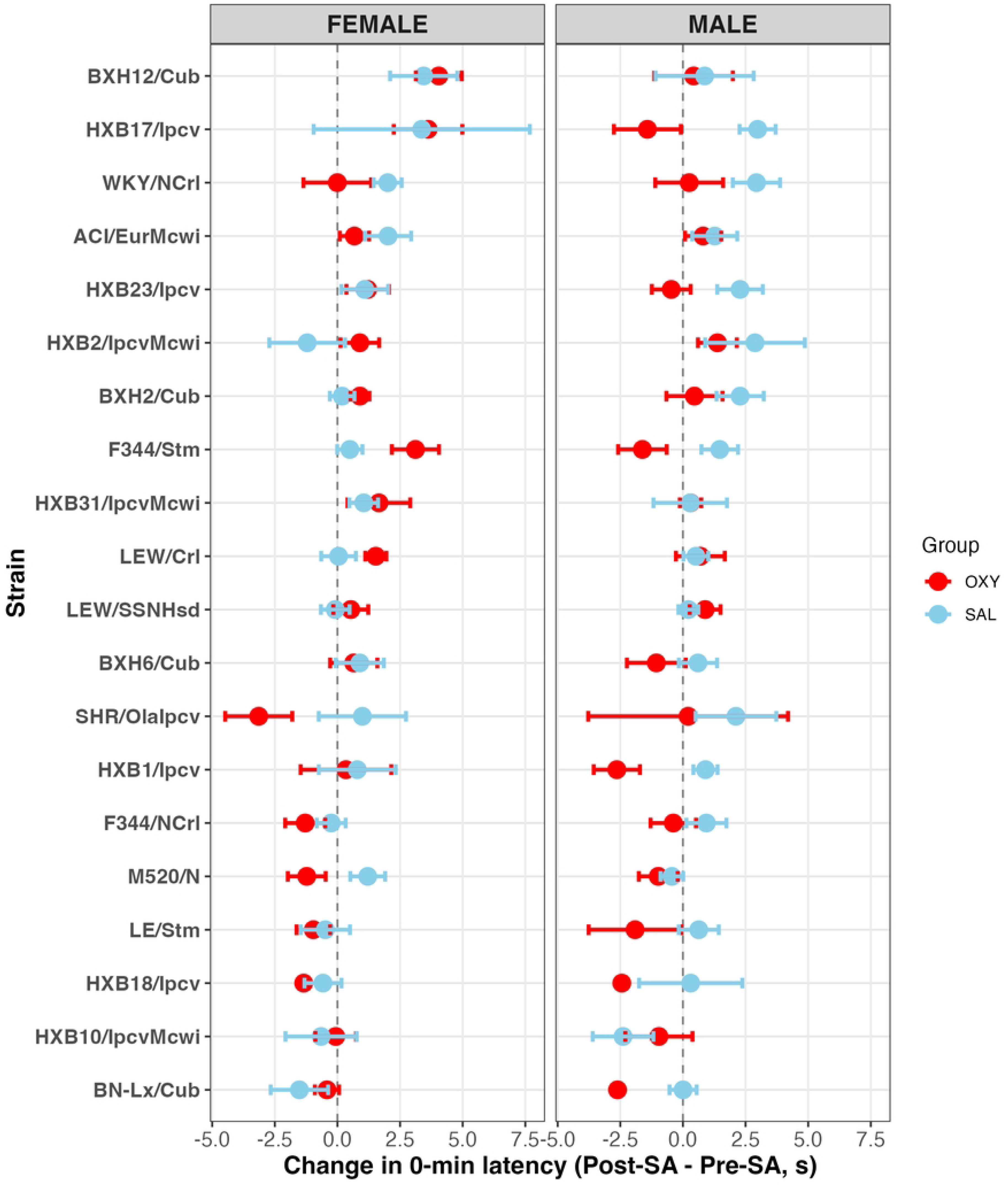
Change in thermal sensitivity following self-administration. Mean change in 0-min tail-withdrawal latency was calculated as Post-SA latency minus Pre-SA latency for each rat and is shown by strain, sex, and SA group. Negative values indicate shorter Post-SA latencies and therefore increased thermal sensitivity after SA. Error bars indicate SEM.

### Oxycodone analgesia before self-administration

The Pre-SA %MPE time course was analyzed using a linear mixed-effects model with strain, sex, timepoint, and their interactions as fixed effects and rat ID as a random intercept. The strain × sex interaction was significant (F₁₉,₅₇₀.₈ = 2.10, p = 0.004), indicating that sex differences in Pre-SA oxycodone analgesia varied across strains. The strain × timepoint interaction was also significant (F₇₆,₂₂₈₈.₂ = 4.30, p < 0.001), indicating that the temporal pattern of Pre-SA analgesia differed across strains. The sex × timepoint interaction and strain × sex × timepoint interaction were not significant.

The strain-specific temporal response patterns, displayed separately by sex, are illustrated in the Pre-SA %MPE heatmap (Fig 2A). Consistent with the significant strain × timepoint interaction, some strains showed rapid, high-magnitude analgesic responses, whereas others showed lower or more limited responses across the assessment period. To provide a simplified example of the strain-level differences apparent in the heatmap, HXB1 and HXB31 are shown as representative high- and low-analgesia strains, averaged across sex (Fig 2B). HXB1 showed a robust early analgesic response, whereas HXB31 showed a substantially lower response across the time course.

**Fig 2.**
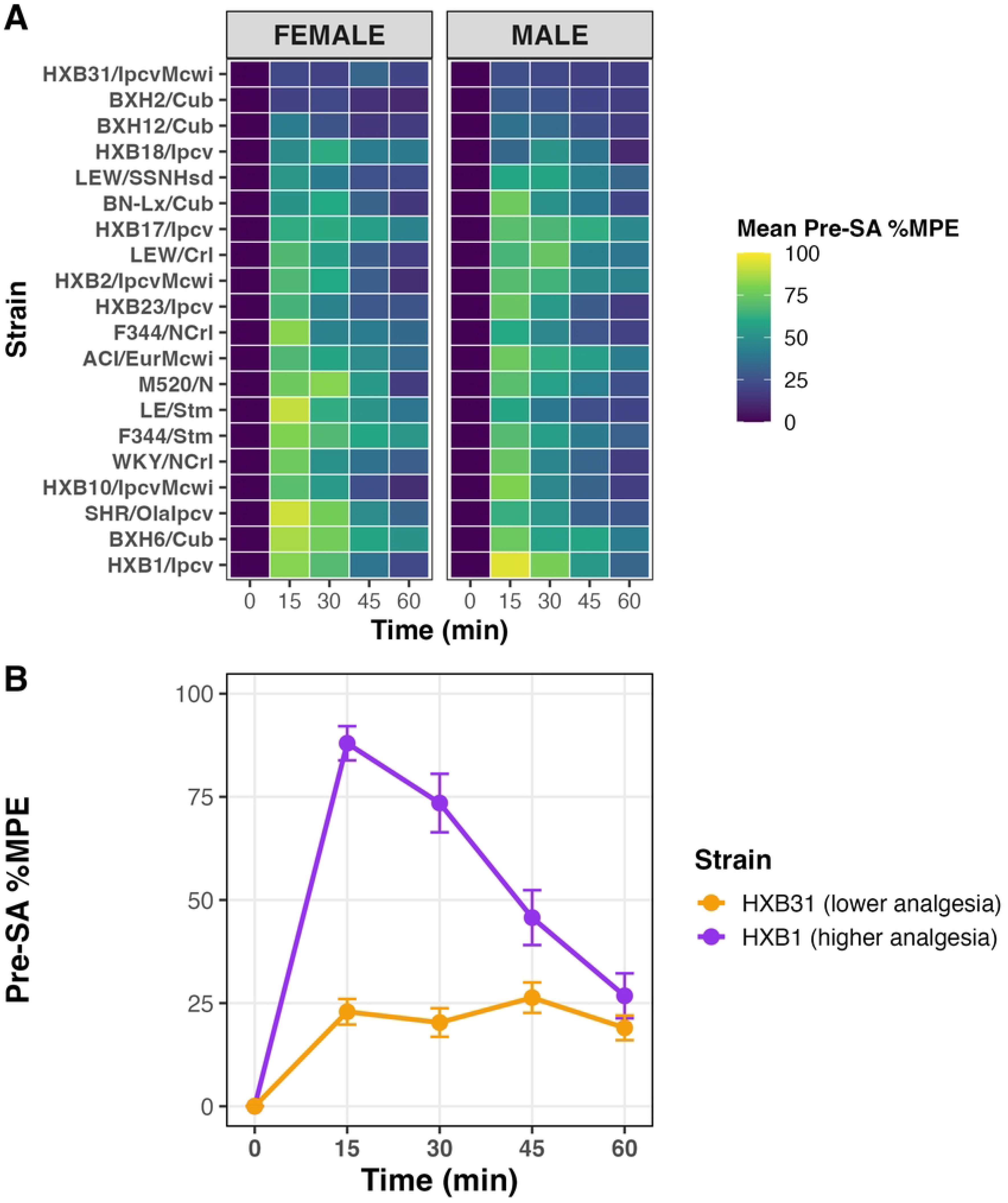
Pre-SA oxycodone analgesia across strains. (A) Heatmap showing mean Pre-SA percent maximum possible effect (%MPE) at 0, 15, 30, 45, and 60 min following acute oxycodone administration, displayed separately for females and males. Each row represents one HRDP strain. Strains are ordered according to mean Pre-SA %MPE at 15 min across both sexes, with lower-response strains at the top and higher-response strains at the bottom. The color scale represents the magnitude of the analgesic response, with higher values indicating greater oxycodone-induced analgesia. (B) Representative Pre-SA %MPE time courses are shown for HXB1 and HXB31, averaged across sex. HXB1 is shown as an example of a strain with a rapid, high-magnitude analgesic response, whereas HXB31 is shown as an example of a strain with lower Pre-SA analgesic responsiveness across the assessment period. Error bars indicate SEM.

To summarize the overall magnitude of Pre-SA oxycodone analgesia, the area under the %MPE–time curve from 0 to 60 min was calculated as the Pre-SA %MPE AUC. Pre-SA %MPE AUC was analyzed using a two-way ANOVA with strain, sex, and their interaction as factors. The strain × sex interaction was significant (F₁₉,₅₇₂ = 2.13, p = 0.004), further indicating that sex differences in overall Pre-SA oxycodone analgesic responsiveness varied across strains (Fig 3). However, Bonferroni-adjusted strain-specific female–male comparisons did not identify any individual strain with a significant sex difference after correction for multiple comparisons.

**Fig 3.**
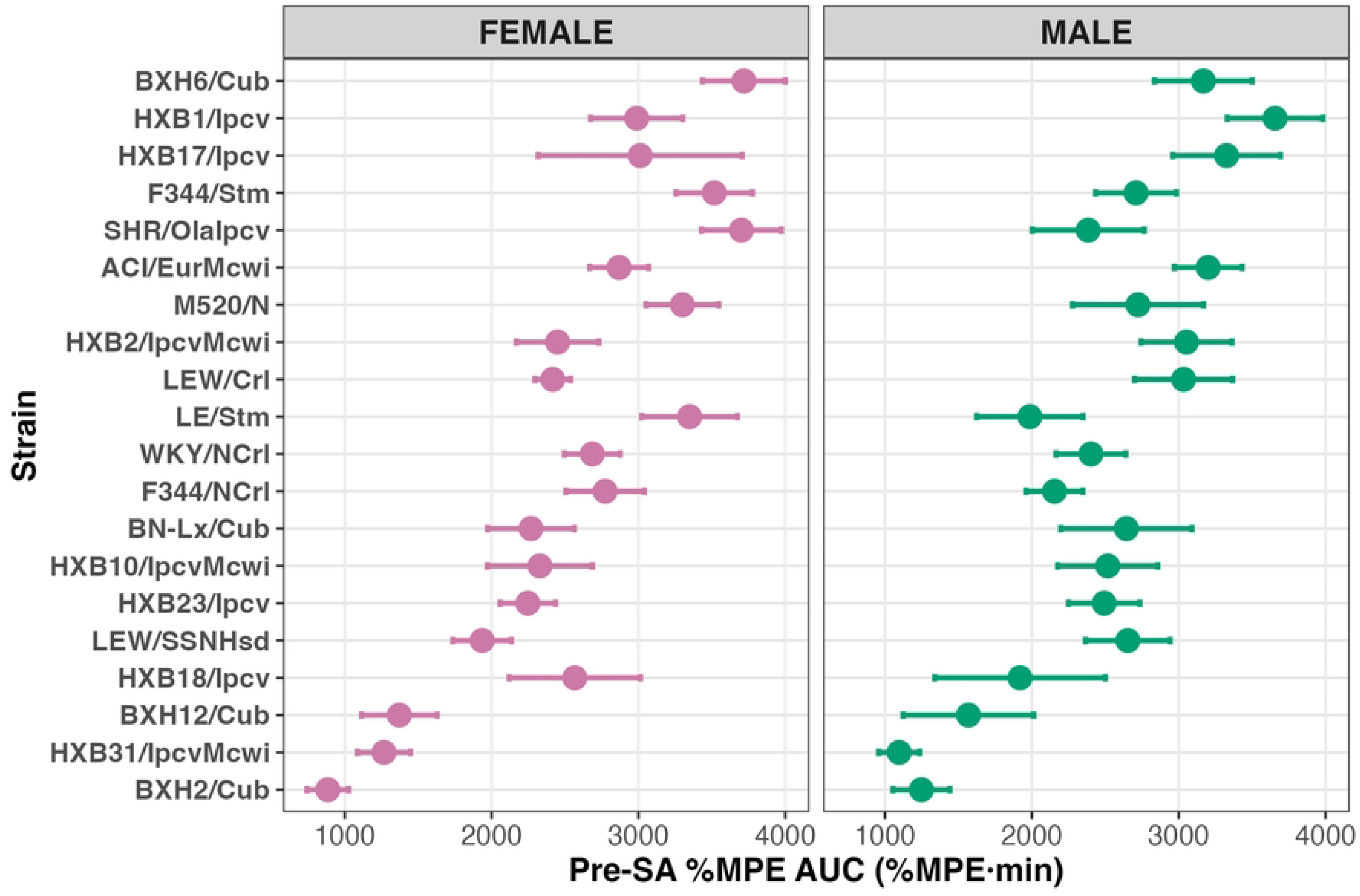
Mean Pre-SA %MPE AUC from 0 to 60 min is shown for each HRDP strain, separately for females and males. Strains are ordered according to overall mean Pre-SA %MPE AUC across both sexes, with lower-response strains at the top and higher-response strains at the bottom. Higher values indicate greater overall oxycodone analgesic responsiveness before SA. Error bars indicate SEM.

### Voluntary oxycodone self-administration produces reduced analgesic responsiveness across genetically diverse rats

Analgesic tolerance was evaluated using complementary between-group and within-subject approaches based on Pre-SA and Post-SA tail-immersion measures. The between-group approach tested whether rats in the Oxy-SA group showed lower analgesic responsiveness during the Post-SA tail-immersion test compared with rats in the Sal-SA group.

The Post-SA %MPE time course was first analyzed using a full linear mixed-effects model that included strain, SA group, timepoint, sex, and their interactions as fixed effects, with rat ID included as a random intercept. Sex and sex-related interaction terms were not significant in the full model. A likelihood-ratio test comparing the full model with a reduced model excluding sex-related terms indicated that sex-related terms did not improve model fit (χ²₂₀₀ = 215.09, p = 0.221). Therefore, the final Post-SA time-course model included strain, SA group, timepoint, and their interactions as fixed effects, with rat ID included as a random intercept.

Significant strain × SA group (F₁₉,₄₆₅.₃ = 1.84, p = 0.0166), strain × timepoint (F₇₆,₁₈₅₉.₆ = 3.20, p < 0.001), and SA group × timepoint interactions (F₄,₁₈₅₉.₇ = 6.50, p < 0.001) were observed in the Post-SA %MPE time-course analysis. The strain × SA group interaction indicated that the difference between the Oxy-SA and Sal-SA groups varied across genetic backgrounds, while the strain × timepoint interaction indicated that the temporal pattern of Post-SA analgesia differed across strains. The SA group × timepoint interaction indicated that the difference between the Oxy-SA and Sal-SA groups depended on time after the acute oxycodone injection.

Bonferroni-adjusted post hoc comparisons following the significant SA group × timepoint interaction showed that rats in the Sal-SA group had greater Post-SA %MPE than rats in the Oxy-SA group at 15 min, p < 0.001; 30 min, p < 0.001; and 45 min, p = 0.011. Post hoc comparisons following the significant strain × SA group interaction showed that, although several strains had nominal group differences before correction, only ACI/EurMcwi and M520/N remained significant after Bonferroni correction. Together, these findings indicate that rats in the Oxy-SA group showed reduced Post-SA analgesic responsiveness during the principal post-injection period, although the magnitude of this difference varied across strains (Fig 4A). To summarize these between-group differences in overall Post-SA analgesic responsiveness, the area under the Post-SA %MPE–time curve from 0 to 60 min was calculated as the Post-SA %MPE AUC. Post-SA %MPE AUC was first analyzed using a full factorial model with strain, sex, SA group, and their interactions as factors. Sex and sex-related interaction terms were not significant in the full model. A likelihood-ratio test comparing the full model with a reduced model excluding sex-related terms indicated that sex-related terms did not improve model fit (χ²₄₀ = 43.35, p = 0.330). Therefore, the final Post-SA %MPE AUC model included strain, SA group, and their interaction as factors. In the final model, the strain × SA group interaction was significant (F₁₉,₄₆₄ = 1.82, p = 0.019), indicating that the difference in overall Post-SA analgesic responsiveness between the Oxy-SA and Sal-SA groups varied across strains. Post hoc comparisons showed that ACI/EurMcwi and M520/N remained significant after Bonferroni correction, with Oxy-SA rats showing lower Post-SA %MPE AUC than Sal-SA rats in both strains. Overall, rats in the Oxy-SA group generally had lower Post-SA %MPE AUC values than rats in the Sal-SA group, providing between-group evidence of reduced oxycodone analgesic responsiveness following oxycodone SA (Fig 4B).

**Fig 4.**
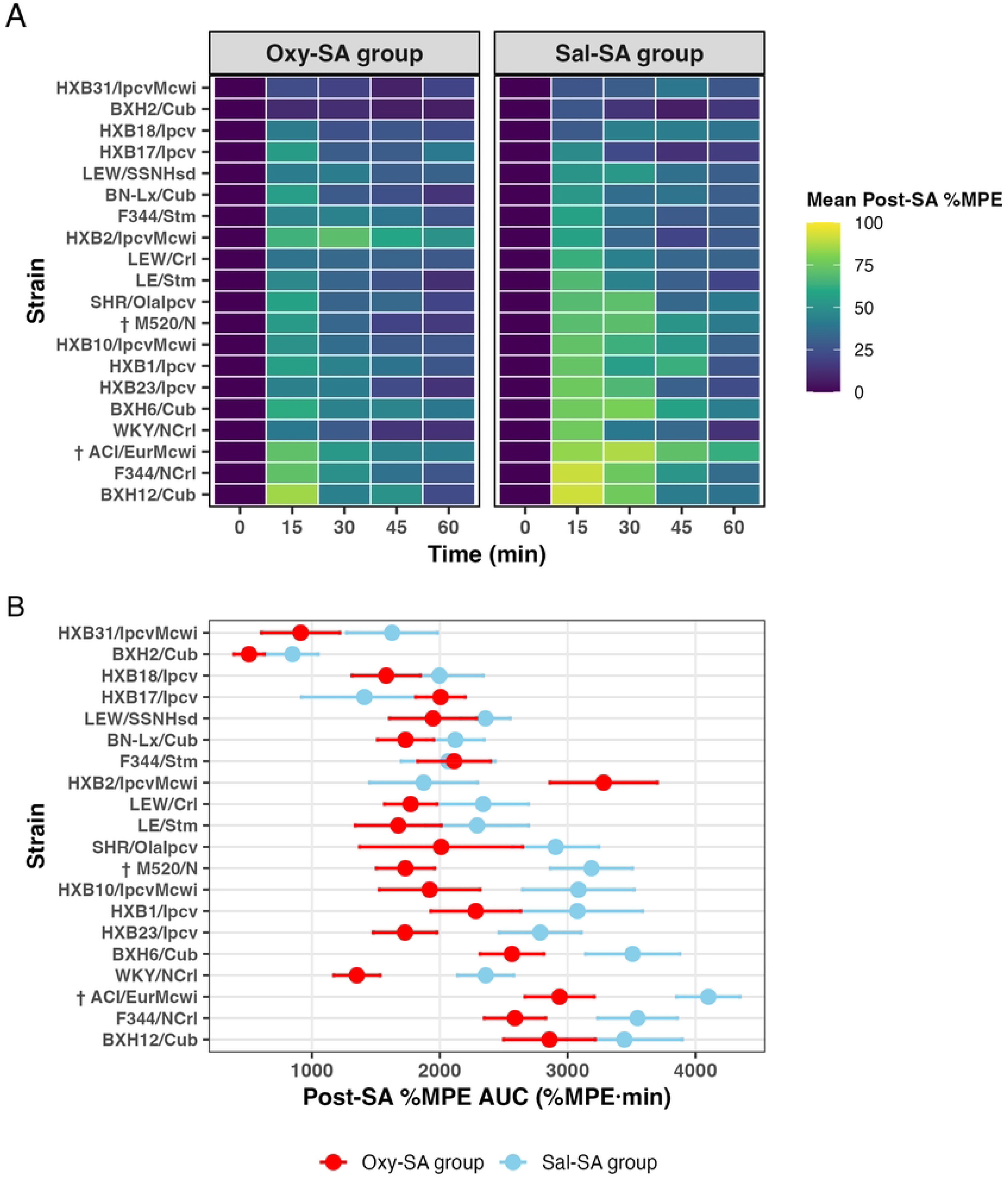
Post-SA oxycodone analgesia following self-administration. (A) Heatmap depicting mean Post-SA percent maximum possible effect (%MPE) across strains in the Oxy-SA and Sal-SA groups at 0, 15, 30, 45, and 60 min during the Post-SA tail-immersion test. Strains are ordered by mean Post-SA %MPE at 15 min in the Sal-SA group, with lower-response strains at the top and higher-response strains at the bottom. Higher %MPE values indicate greater oxycodone analgesia. † indicates strains that remained significant after Bonferroni-corrected post hoc comparisons for the strain × SA group interaction. Overall, rats in the Oxy-SA group showed lower Post-SA %MPE than rats in the Sal-SA group, providing between-group evidence of reduced oxycodone analgesic responsiveness following oxycodone SA. (B) Mean Post-SA %MPE AUC from 0 to 60 min is shown for each HRDP strain in the Oxy-SA and Sal-SA groups. Strains are ordered according to mean Post-SA %MPE AUC in the Sal-SA group, with lower-response strains at the top and higher-response strains at the bottom. Lower Post-SA %MPE AUC values indicate reduced overall oxycodone analgesic responsiveness after SA. Error bars indicate SEM. † indicates strains with a significant Bonferroni-corrected Oxy-SA versus Sal-SA comparison for Post-SA %MPE AUC.

To examine within-subject changes in analgesic responsiveness in the Oxy-SA group, Pre-SA and Post-SA %MPE time courses were compared using a linear mixed-effects model with phase, strain, timepoint, and their interactions as fixed effects and rat ID as a random intercept. Significant phase × strain (F₁₉,₂₇₃₄.₉ = 3.94, p < 0.001) and phase × timepoint interactions (F₄,₂₄₉₉.₈ = 10.44, p < 0.001) indicated that Pre-SA-to-Post-SA differences varied across strains and post-injection timepoints. The phase × strain × timepoint interaction was not significant. Bonferroni-adjusted post hoc comparisons identified significantly lower Post-SA %MPE at one or more timepoints in LEW/Crl, WKY/NCrl, HXB23/Ipcv, M520/N, SHR/OlaIpcv, BXH6/Cub, LE/Stm, and HXB1/Ipcv, a pattern consistent with analgesic tolerance with no significant reductions detected in the corresponding Sal-SA strain profiles. BXH12/Cub showed significantly greater Post-SA %MPE at selected timepoints, with a similar increase observed in the Sal-SA group (S2 Fig). These results demonstrate strain-dependent Pre-SA-to-Post-SA changes in analgesic responsiveness (Fig 5).

**Fig 5.**
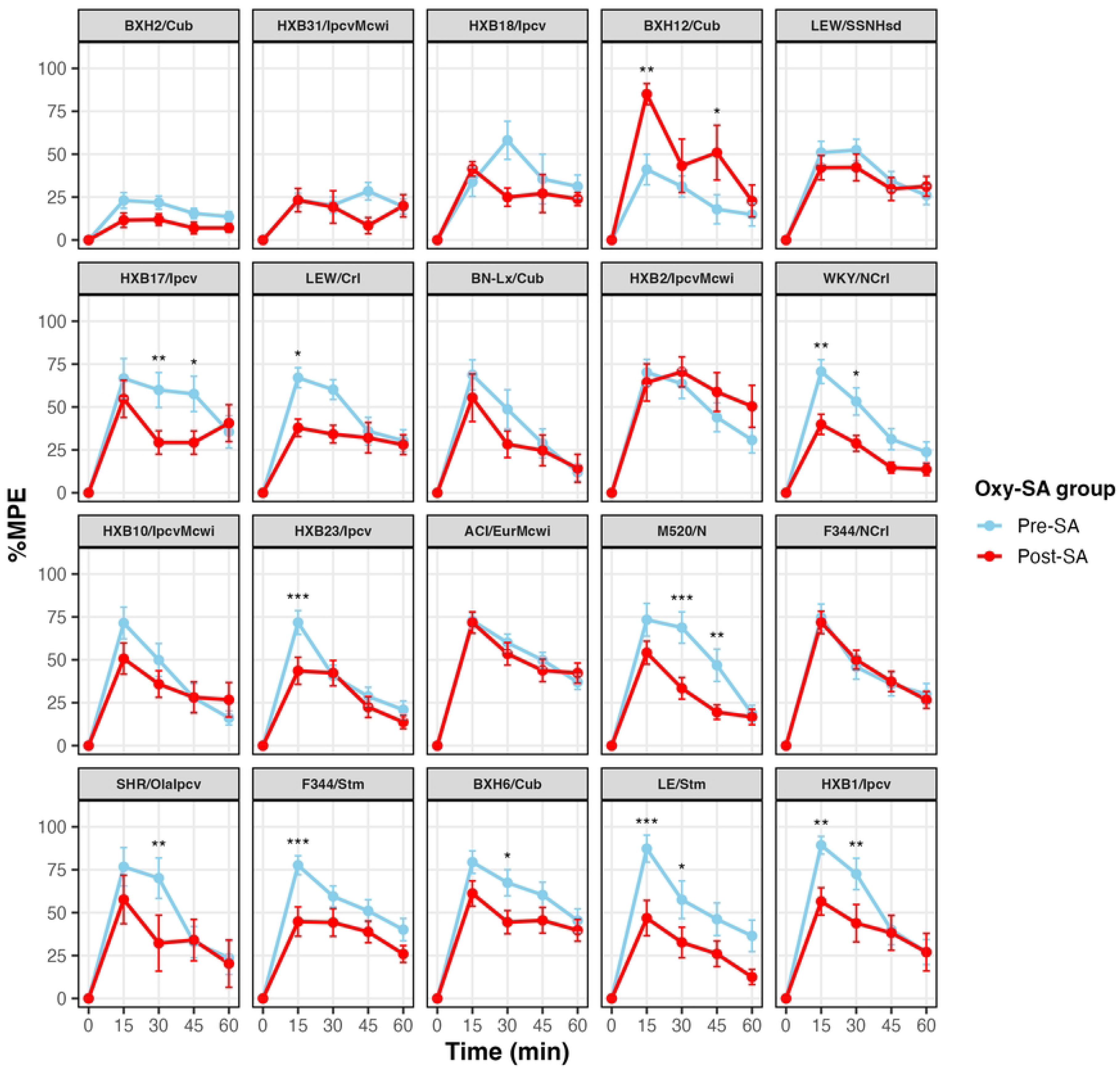
Strain-specific Pre-SA-to-Post-SA changes in oxycodone analgesia in the Oxy-SA group. Pre-SA and Post-SA oxycodone analgesia time courses are shown as mean %MPE ± SEM for each strain. Higher %MPE values indicate greater oxycodone analgesia. Asterisks indicate significant Pre-SA versus Post-SA comparisons at individual post-injection timepoints, with Bonferroni adjustment across the four post-injection timepoints within each strain: *p < 0.05, **p < 0.01, and ***p < 0.001.

Analgesic tolerance was also quantified using a single within-subject summary score, the Tolerance AUC difference, calculated for each rat as Post-SA %MPE AUC minus Pre-SA %MPE AUC. This score was used as the tolerance phenotype for subsequent heritability and intake-correlation analyses. Negative values indicated reduced analgesic responsiveness from Pre-SA to Post-SA in either group. Tolerance AUC difference was analyzed using a three-way factorial ANOVA with strain, sex, SA group, and their interactions as factors. Significant main effects were observed for strain (*F*₁₉,₄₂₂ = 3.47, *p* < 0.001), sex (*F*₁,₄₂₂ = 4.64, *p* = 0.032), and SA group (F₁,₄₂₂ = 8.91, p = 0.003). Females had more negative Tolerance AUC difference values than males overall (p = 0.032). None of the two-way or three-way interactions were significant. The significant strain effect indicated that Pre-SA-to-Post-SA change scores differed among strains overall; however, the nonsignificant strain × SA group interaction indicated that the difference between the Oxy-SA and Sal-SA groups did not vary significantly across strains. Across strains and sexes, rats in the Oxy-SA group had more negative Tolerance AUC difference values than rats in the Sal-SA group, supporting a greater reduction in oxycodone analgesic responsiveness associated with oxycodone SA and consistent with analgesic tolerance (Fig 6).

**Fig 6.**
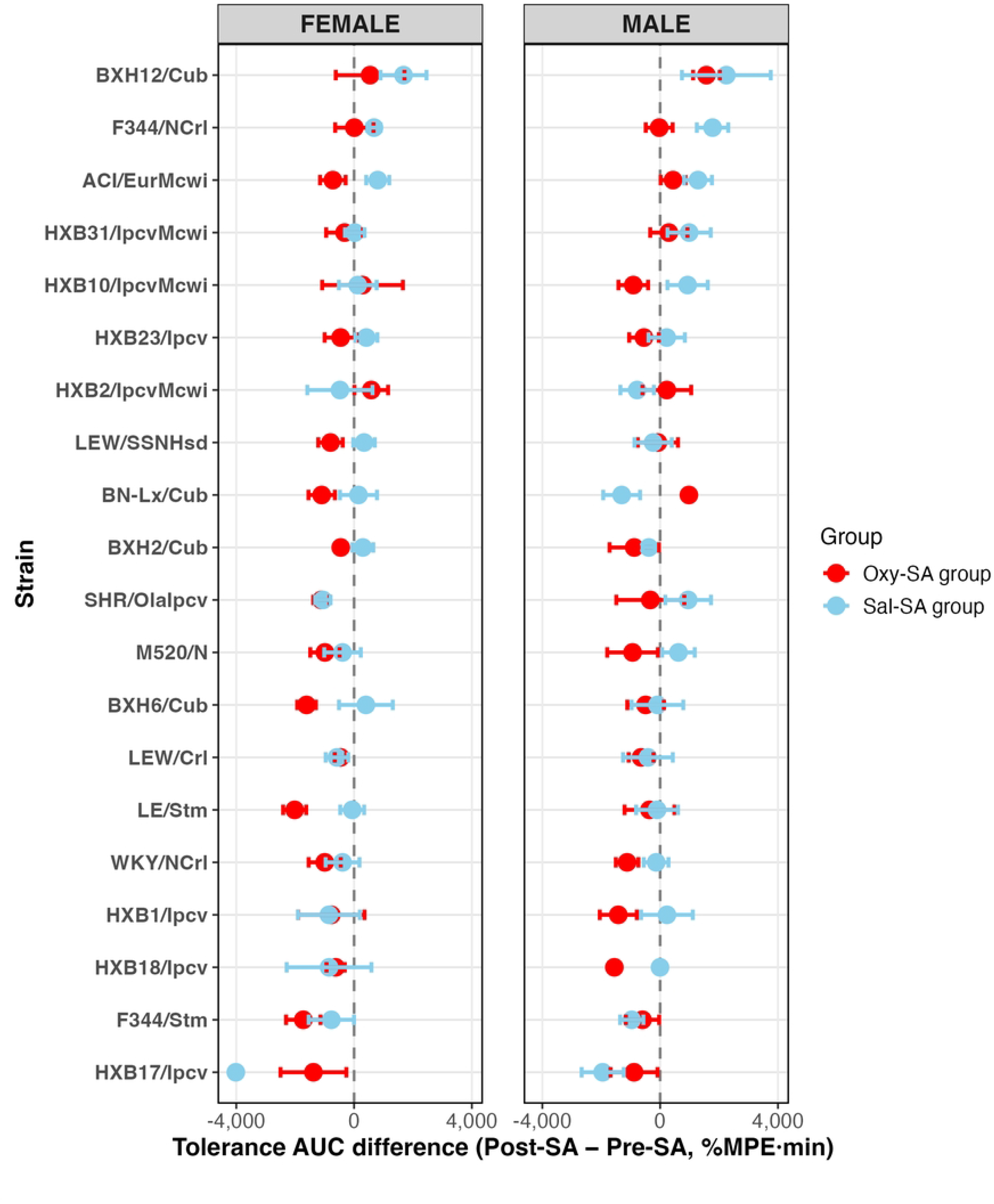
Within-subject analgesic tolerance following self-administration. Tolerance AUC difference was calculated as Post-SA %MPE AUC minus Pre-SA %MPE AUC for each rat. Negative values indicate reduced analgesic responsiveness from Pre-SA to Post-SA. The more negative values observed in the Oxy-SA group relative to the Sal-SA group support analgesic tolerance associated with oxycodone SA. Error bars indicate SEM.

### Relationship between total oxycodone intake and tolerance across genetically diverse rat strains

To evaluate the relationship between oxycodone intake and analgesic tolerance, we calculated correlations between total oxycodone intake (mg/kg) and tolerance AUC difference, defined as Post-SA %MPE AUC minus Pre-SA %MPE AUC. Total oxycodone intake varied substantially across strains (S3 Fig). At the strain level, mean tolerance AUC difference was not associated with mean total oxycodone intake among rats in the Oxy-SA group (r = −0.06, p = 0.79; Fig 7), indicating that strains with greater average oxycodone intake did not necessarily show greater analgesic tolerance. A post hoc sensitivity analysis at 15 min, when analgesia was highest and tolerance-like reductions were most visible, also showed no association between strain-mean total oxycodone intake and the Pre-SA-to-Post-SA change in %MPE (r = 0.02, p = 0.945; S4 Fig).

**Fig 7.**
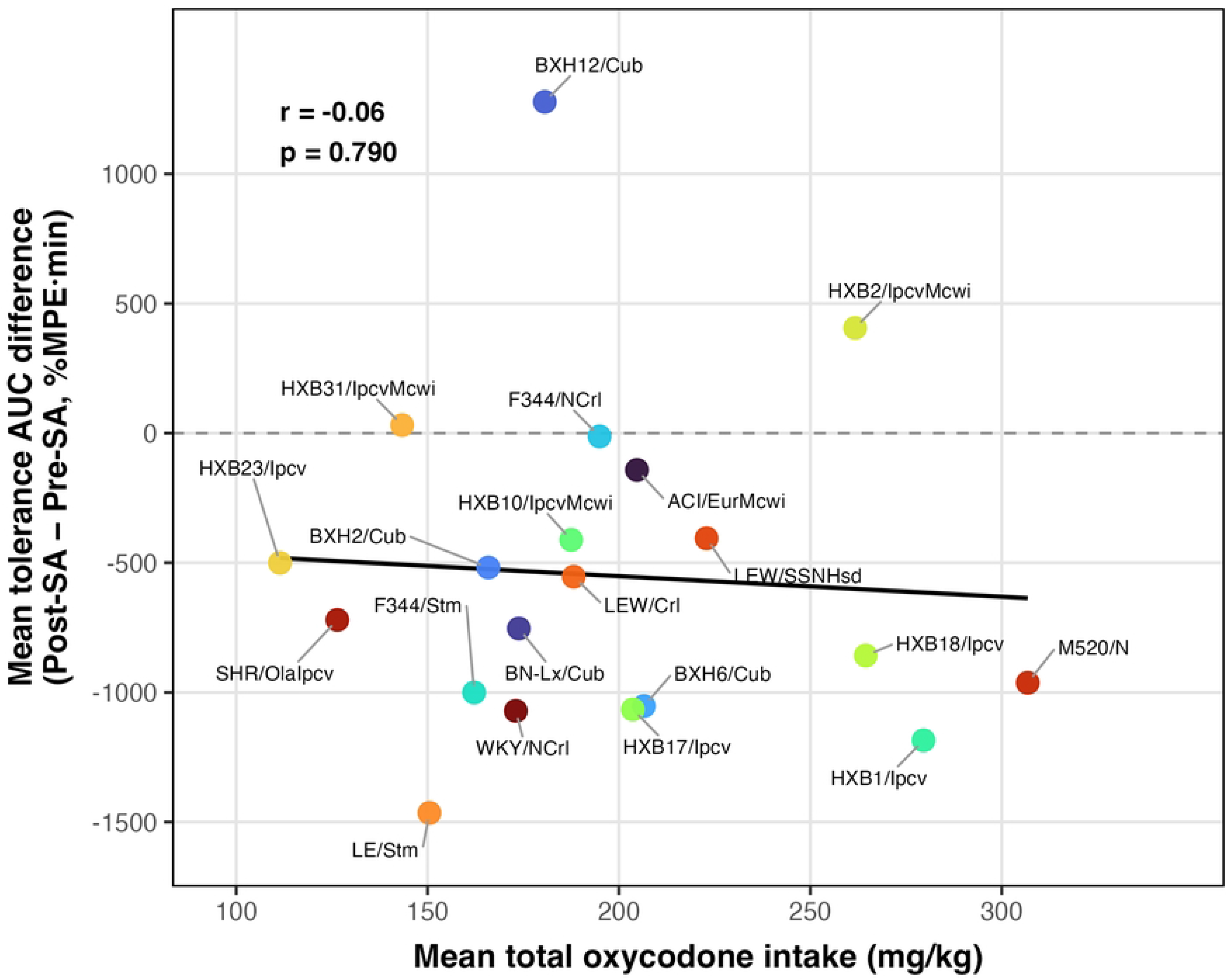
Strain-level relationship between total oxycodone intake and tolerance AUC difference. Each point represents one HRDP strain, with mean total oxycodone intake among Oxy-SA rats on the x-axis and mean tolerance AUC difference on the y-axis. Tolerance AUC difference was calculated as Post-SA %MPE AUC − Pre-SA %MPE AUC. The fitted regression line showed no meaningful association between strain-average oxycodone intake and mean tolerance AUC difference (r = −0.06, p = 0.79).

We next examined whether total oxycodone intake and tolerance AUC difference were associated among individual rats within each strain. Within-strain correlations were generally small and varied in direction (Fig 8). Only one nominally significant association was observed across the strain panel, in BXH6/Cub (r = −0.59, p = 0.026). Collectively, these findings suggest that variation in analgesic tolerance in this paradigm is not well explained by total oxycodone intake and that oxycodone intake and analgesic tolerance may represent at least partly separable traits.

**Fig 8.**
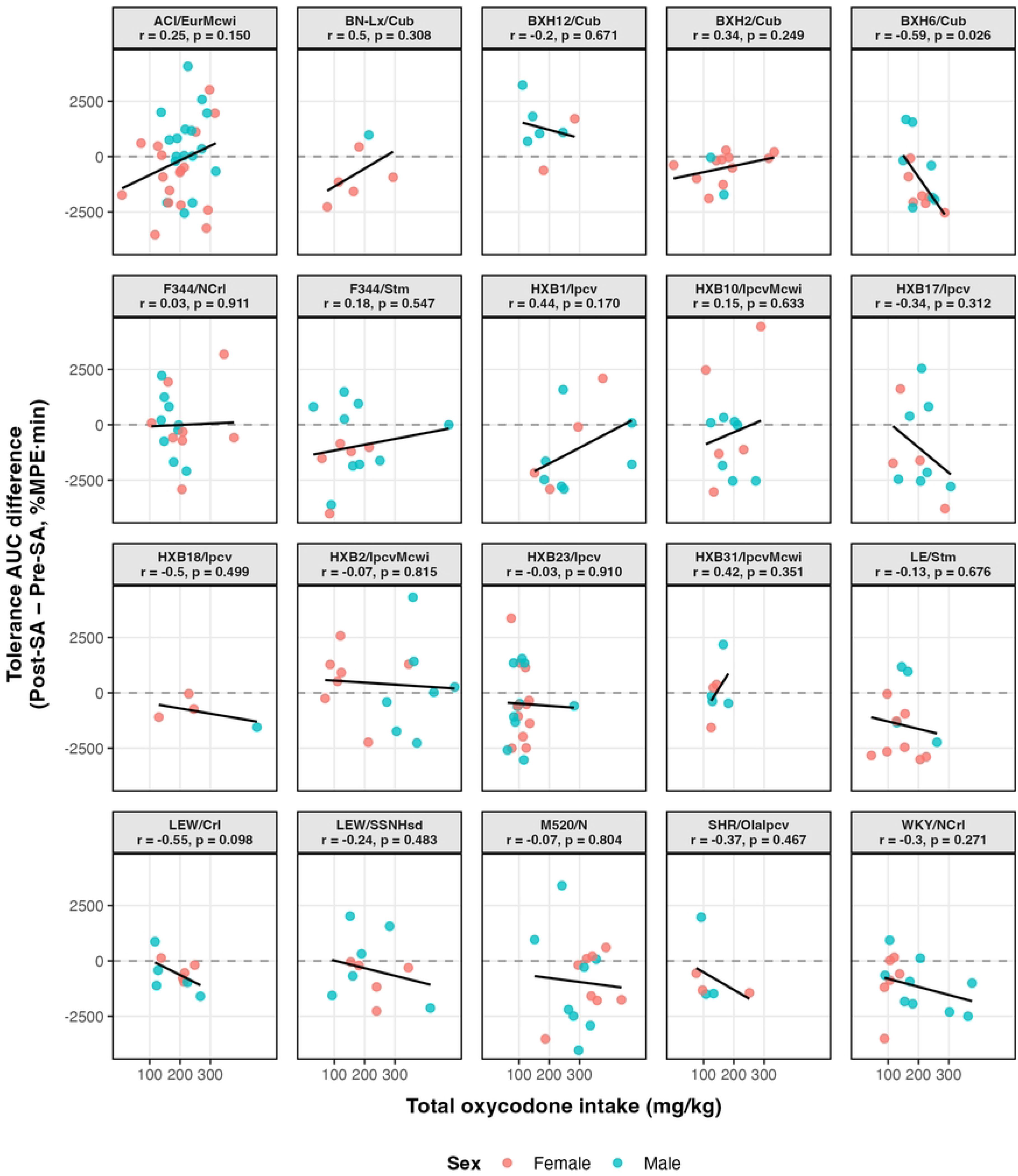
Within-strain relationships between total oxycodone intake and tolerance AUC difference. Each point represents an individual oxycodone self-administering rat and is colored by sex. Separate panels show the association between total oxycodone intake and tolerance AUC difference within each HRDP strain, with a fitted linear regression line displayed for each strain. Tolerance AUC difference was calculated as Post-SA %MPE AUC − Pre-SA %MPE AUC.

### Heritability estimate of behavioral phenotypes

Heritability estimates indicated that genetic background accounted for a moderate share of the variance in oxycodone analgesia and thermal sensitivity (H² ≈ 0.28–0.40; Table 3), both before and after SA. In contrast, analgesic tolerance and change in thermal sensitivity showed lower heritability (H² ≈ 0.18–0.27; Table 3), suggesting that these adaptation-related traits may be less dependent on genetic factors in this panel.

**Table 3.** Heritability estimates for opioid-related behavioral phenotypes in HRDP rat strains. . Broad-sense heritability estimates (H²) are shown for analgesia, tolerance, and thermal sensitivity phenotypes in the Oxy-SA and Sal-SA groups.

| PHENOTYPES | OXY | SAL |
| --- | --- | --- |
| Pre-SA thermal sensitivity | 0.32 |  |
| Post-SA thermal sensitivity | 0.34 | 0.32 |
| Change in thermal sensitivity | 0.22 | 0.21 |
| Pre-SA analgesia | 0.28 |  |
| Post-SA analgesia | 0.34 | 0.40 |
| Analgesic tolerance | 0.18 | 0.27 |

Broad-sense heritability ( $H^2$ ) estimates are shown for each phenotype. OXY, oxycodone self-administration group; SAL, saline self-administration group; HRDP, Hybrid Rat Diversity Panel; Pre-SA, pre-self-administration; Post-SA, post-self-administration.

## Discussion

This study examined how genetic background and sex influence oxycodone analgesia, analgesic tolerance, and OIH following voluntary oxycodone (or saline) SA across 20 inbred strains from the HRDP. By combining Pre- and Post-SA tail-immersion testing with strain-level analyses, heritability estimation, and correlations between total oxycodone intake and analgesic tolerance, we identified several major findings. First, baseline thermal sensitivity and initial oxycodone analgesia varied substantially across strains, indicating strong genetic regulation of pain sensitivity and opioid analgesic efficacy. Second, voluntary oxycodone intake reduced analgesic efficacy in several strains, consistent with the development of analgesic tolerance. Third, oxycodone SA produced hyperalgesia, an effect most evident in males. Finally, total oxycodone intake was not associated with tolerance at the strain level, suggesting that the biological mechanisms governing oxycodone consumption and those governing analgesic tolerance are at least partly dissociable in this model. A novel aspect of this study is the use of voluntary SA rather than experimenter-controlled dosing. Because oxycodone exposure was determined by the animals’ own intake, this approach may better capture individual variability in opioid-taking behavior. This translational relevance is supported by prior rodent studies using opioid SA to model voluntary opioid intake and related behavioral adaptations [11,21–26].

### Thermal sensitivity and opioid-induced hyperalgesia

We first observed substantial strain-dependent variation in baseline thermal sensitivity, indicating that genetic background contributes strongly to basal nociceptive sensitivity in the HRDP. This finding is consistent with prior work showing that pain sensitivity differs across genetically distinct rodent populations [7,13,27,28]. After SA, Oxy-SA rats showed a greater reduction in 0-min tail-withdrawal latency than Sal-SA rats, an effect primarily observed in males, indicating increased thermal sensitivity consistent with OIH. This finding aligns with Liu et al., who reported hyperalgesia following repeated opioid exposure in rodents [29], and with broader evidence that repeated opioid exposure can enhance pain sensitivity in both clinical and preclinical settings [30–33]. Although Liu et al. tested for sex differences in OIH, no significant sex effect was observed following experimenter-administered morphine. We observed greater OIH in males suggesting that the opioid type and route of administration may play a role in the sex-dependent expression of OIH. Although change in thermal sensitivity differed across strains overall, this strain effect was not specific to oxycodone SA. Thus, the present findings suggest that opioid-induced hyperalgesia under voluntary intake conditions may be more strongly influenced by sex than by genetic background, and that the route of opioid exposure (voluntary SA versus experimenter-administered injection) may itself shape whether sex differences emerge.

### Oxycodone analgesia and analgesic tolerance

Oxycodone analgesia tested before SA varied markedly across HRDP strains, indicating that genetic background strongly influences initial opioid analgesic efficacy. This was evident in both the %MPE time course analysis and the Pre-SA analgesia AUC, where some strains showed rapid and robust increases in analgesia, whereas others showed weaker or shorter-lasting responses. These findings are consistent with prior rodent studies showing that opioid analgesia differs substantially across genetic backgrounds [7,12,18].

Following SA, rats in the Oxy-SA group showed reduced Post-SA analgesic responses compared with rats in the Sal-SA group, providing between-group evidence of reduced oxycodone analgesic responsiveness. This reduction was evident in the Post-SA %MPE time course and in the lower Post-SA %MPE AUC observed in the Oxy-SA group. The strain-specific Pre-SA to Post-SA time courses showed that several strains displayed a tolerance-like pattern, characterized by reduced Post-SA analgesia compared with Pre-SA analgesia at multiple post-injection timepoints. This pattern was further supported by within-subject AUC difference score comparisons, where rats in the Oxy-SA group showed greater shifts toward negative AUC difference scores than rats in the Sal-SA group. Together, these findings are consistent with prior preclinical work showing that repeated opioid exposure produces analgesic tolerance in rodents [12,33–38], as well as human clinical and experimental studies showing reduced opioid analgesic efficacy or increased opioid requirement following repeated opioid exposure [39–41].

A notable feature of the present study was that tolerance-related responses were not expressed uniformly across strains. Several strains in the Oxy-SA group showed tolerance-like patterns characterized by significantly lower Post-SA analgesia than Pre-SA analgesia at one or more post-injection timepoints (e.g., LEW/Crl, WKY/NCrl, HXB23/Ipcv, M520/N, SHR/OlaIpcv, BXH6/Cub, LE/Stm, and HXB1/Ipcv). In contrast, BXH12/Cub showed enhanced Post-SA analgesia in both Oxy-SA and Sal-SA groups, suggesting that this increase was not specific to oxycodone SA. Other strains, such as LEW/SSNHsd, showed relatively stable analgesic responses across testing sessions. Thus, voluntary oxycodone SA was associated with an overall reduction in analgesic responsiveness, although the magnitude and timing of this change varied across strains. This heterogeneity highlights the complexity of tolerance-like adaptations in genetically diverse populations and is consistent with prior studies showing that genetic background can influence opioid analgesia and tolerance. For example, comparisons among Wistar-Kyoto (WKY), Sprague Dawley, and spontaneously hypertensive (SHR) rats have demonstrated strain-related differences in morphine sensitivity (SHR > WKY) and in the development of tolerance (WKY > SHR) following repeated experimenter-delivered morphine exposure [42]. Interestingly, the WKY/NCrl and SHR/OlaIpcv strains evaluated in our studies showed relatively similar analgesic and tolerance profiles suggesting that opioid type and repeated exposure paradigm may be important factors. Regardless, these findings support the view that genetic background is an important determinant of both the initial analgesic response to opioids and the subsequent development of tolerance.

This heterogeneity is also consistent with Kuhn et al., who used a heroin use disorder model in more than 900 male and female outbred heterogeneous stock rats to identify distinct vulnerable, intermediate, and resilient behavioral profiles [25]. In that study, rats underwent long-access heroin SA and several heroin-related behavioral tests, and nonlinear clustering was used to classify animals based on OUD-like behavioral traits. Although analgesic tolerance was not directly assessed, their findings support the broader concept that opioid-related adaptations are multidimensional and heterogeneous rather than uniform across individuals. The HRDP extends this idea by allowing analgesic adaptation to be examined across a broad range of genetic backgrounds.

A key question in this study was whether the magnitude of analgesic tolerance was explained by the amount of oxycodone consumed during SA. Previous rodent work has shown that opioid dosing protocols and cumulative exposure can strongly influence the development of tolerance, with continuous infusion generally producing greater tolerance than acute or intermittent administration even at comparable daily doses, and with differences across opioid agonists such as etorphine, methadone, oxycodone, and hydrocodone [43]. Based on this literature, we hypothesized that rats and strains with higher voluntarily self-administered oxycodone intake would exhibit greater analgesic tolerance. Mean Tolerance AUC Difference scores showed no meaningful relationship with mean total oxycodone intake when analyzed at the strain-level, indicating that strains self-administering more oxycodone did not necessarily develop analgesic tolerance. We next examined whether intake and tolerance were related within individual strains.

Similarly, within-strain correlations were generally small and inconsistent in direction indicating that individual rats with higher oxycodone intake did not consistently show greater tolerance within their strain. To our knowledge, this study provides one of the first direct tests of whether voluntarily self-administered oxycodone intake predicts the magnitude of analgesic tolerance across a genetically diverse rat panel. This dissociation between intake and tolerance challenges the assumption that greater opioid consumption produces greater analgesic tolerance. Although drug exposure is clearly necessary for tolerance to develop, the present findings suggest that total exposure alone is not sufficient to explain individual- or strain-level differences in tolerance magnitude.

One limitation is that total oxycodone intake was analyzed as a cumulative exposure measure across the 20-day SA protocol, including both acquisition and escalation phases. Although this measure captures overall voluntary drug exposure, it does not distinguish whether tolerance is more strongly related to specific phases or patterns of intake, such as early acquisition, long-access escalation, within-session distribution of infusions, or burst-like intake. However, the absence of an association between intake and tolerance was not limited to the AUC-based measure. At the 15-min post-injection timepoint, when the analgesic response was greatest and tolerance-like reductions were most apparent across strains, mean total oxycodone intake was not associated with the strain-mean Pre-SA-to-Post-SA change in %MPE (r = 0.02, p = 0.945; S4 Fig). This timepoint-specific analysis supports the AUC-based findings and indicates that the lack of association was unlikely to result from averaging analgesic responses across the 0–60-min time course. Together, these results suggest that cumulative oxycodone intake alone does not explain strain differences in the loss of analgesic efficacy and highlight the potential contributions of genetic and biological factors to analgesic tolerance.

### Heritability of oxycodone-related pain phenotypes

Broad-sense heritability estimates indicated that genetic background makes a substantial contribution to several oxycodone-related pain phenotypes in the HRDP panel. Both Pre-SA and Post-SA oxycodone analgesia showed moderate heritability, with H² values ranging from approximately 0.28 to 0.40 across SA groups. Similar H² values were observed for Pre-SA and Post-SA thermal sensitivity (H² ≈ 0.32–0.34), indicating that genetic factors explain a sizeable fraction of the variance in baseline nociception and in pain sensitivity after chronic oxycodone exposure. These results are consistent with prior rodent studies reporting moderate-to-high heritability for baseline pain sensitivity [44,45], strain-dependent variation in opioid-induced analgesia [7,9,19], and human twin data demonstrating that genetic factors account for roughly one quarter to one third of the variability in acute opioid analgesic response and pain sensitivity [46,47]. Together, these findings support the idea that acute opioid sensitivity and baseline pain traits are shaped by inherited differences in pain and opioid systems.

In contrast, heritability estimates for opioid-induced adaptations such as tolerance and hyperalgesia were lower than those for acute analgesia but still clearly non-zero. Analgesic tolerance exhibited H² values of approximately 0.18–0.27 and change in thermal sensitivity showed H² values of approximately 0.21–0.22, indicating a measurable but more modest genetic contribution to these phenotypes. This pattern suggests that environmental influences and possibly gene-by-environment interactions (e.g., strain-dependent variability in response to oxycodone exposure) may account for a larger proportion of the observed variability. In addition, both tolerance and change in thermal sensitivity are difference-score measures, which can combine measurement variability across timepoints and reduce heritability estimates even when underlying genetic effects are present. Importantly, the lower heritability estimates observed here do not mean that opioid tolerance is not genetically influenced. Consistent with this interpretation, Kest et al. surveyed morphine tolerance across multiple inbred mouse strains and observed robust strain-dependent variation in tolerance development, supporting a genetic contribution to opioid tolerance in rodent models [20].

In humans, Gaddis et al. reported SNP-based heritability estimates ranging from h² = 0.11 to 0.18 across opioid-addiction phenotypes [15]. Although the present HRDP broad-sense estimates, human twin estimates, and SNP-based estimates all support a genetic contribution, it is difficult to directly compare these estimates due to different study designs and capturing different components of genetic variation. In addition, the Gaddis et al. estimates reflect addiction liability rather than tolerance specifically. Nevertheless, this finding remains relevant because tolerance may contribute to dose escalation and, ultimately, dependence [48]. Viewed together, the present HRDP results and existing rodent and human studies suggest that genetics plays a role in baseline nociception and acute analgesia, and a smaller but still detectable role in opioid-induced tolerance, with environmental exposures, individual drug-use patterns, and gene-by-environment interactions likely contributing additional influence on these adaptations.

## Conclusion

In conclusion, genetic background contributed to variation in baseline thermal sensitivity and oxycodone analgesia across HRDP strains. Voluntary oxycodone SA produced reduced analgesic responsiveness consistent with tolerance and a sex-dependent increase in thermal sensitivity that was most evident in males. However, total oxycodone intake was not associated with tolerance, suggesting that drug consumption and loss of analgesic efficacy are partly dissociable traits. These findings highlight individual variability in opioid responses and provide a foundation for future studies investigating the biological mechanisms underlying vulnerability or resistance to opioid tolerance.

## Acknowledgments

The authors thank Melinda Dwinell PhD (Medical College of Wisconsin) for providing many of the inbred strains studied in this article. We gratefully acknowledge the contributions of Safa Vaseemuddin and Cove Andrews, whose dedication and technical assistance were instrumental in the execution and completion of these studies. We also appreciate the thoughtful discussion about analyses with Ms. Alanna Mayberry.

## Supporting information

**S1 Fig Baseline tail-withdrawal latency at the Pre-SA 0-min timepoint.** Mean tail-withdrawal latency values (± SEM) are shown for each HRDP rat strain, separately for females and males. Strains are ordered according to their overall mean baseline latency across both sexes. Longer withdrawal latencies indicate lower thermal sensitivity, whereas shorter latencies indicate greater thermal sensitivity.

**S2 Fig. Pre-SA and Post-SA oxycodone analgesia in saline self-administering rats.** Mean %MPE ± SEM is shown for each strain before and after saline self-administration. Asterisks indicate significant Pre-SA versus Post-SA comparisons at individual post-injection timepoints, with Bonferroni adjustment applied across the four post-injection timepoints within each strain: *p < 0.05, **p < 0.01, and ***p < 0.001.

**S3 Fig Mean total oxycodone intake by HRDP strain.** Mean total oxycodone dose (mg/kg) self-administered by each HRDP rat strain is displayed, with error bars indicating SEM. Strains are ranked from lowest to highest mean intake.

**S4 Fig. Strain-level relationship between total oxycodone intake and the 15-min tolerance difference.** Each point represents one HRDP strain. Mean total oxycodone intake among Oxy-SA rats is plotted against the mean 15-min tolerance difference, calculated as Post-SA %MPE minus Pre-SA %MPE. Negative values indicate lower Post-SA analgesic responsiveness. The 15-min timepoint was selected because analgesia was highest and tolerance-like reductions were most apparent. No association was observed between total oxycodone intake and the 15-min tolerance difference (r= 0.02, p = 0.945). The line represents the fitted linear regression.

**S1 Data.** Animal identifiers and raw rat-level tail-immersion, and total reinforced infusions and total oxycodone intake across the self-administration protocol.

## References

1. Chou R, Turner JA, Devine EB, Hansen RN, Sullivan SD, Blazina I, et al. The Effectiveness and Risks of Long-Term Opioid Therapy for Chronic Pain: A Systematic Review for a National Institutes of Health Pathways to Prevention Workshop. Ann Intern Med. 2015;162: 276–286. doi:10.7326/M14-2559

2. Dowell D, Ragan KR, Jones CM, Baldwin GT, Chou R. CDC Clinical Practice Guideline for Prescribing Opioids for Pain — United States, 2022. MMWR Recomm Rep. 2022;71: 1–95. doi:10.15585/mmwr.rr7103a1

3. Mercadante S, Arcuri E, Santoni A. Opioid-Induced Tolerance and Hyperalgesia. CNS Drugs. 2019;33: 943–955. doi:10.1007/s40263-019-00660-0

4. Silverman SM. Opioid Induced Hyperalgesia: Clinical Implications for the Pain Practitioner. Pain Physician. 2009;3;12: 679–684. doi:10.36076/ppj.2009/12/679

5. Doverty M, White JM, Somogyi AA, Bochner F, Ali R, Ling W. Hyperalgesic responses in methadone maintenance patients. Pain. 2001;90: 91–96. doi:10.1016/S0304-3959(00)00391-2

6. Sibille KT, Kindler LL, Glover TL, Gonzalez RD, Staud R, Riley JL, et al. Individual Differences in Morphine and Butorphanol Analgesia: A Laboratory Pain Study. Pain Med. 2011;12: 1076–1085. doi:10.1111/j.1526-4637.2011.01157.x

7. Duffy EP, Ward JO, Hale LH, Brown KT, Kwilasz AJ, Saba LM, et al. Genetic background and sex influence somatosensory sensitivity and oxycodone analgesia in the Hybrid Rat Diversity Panel. Genes Brain Behav. 2024;23: e12894. doi:10.1111/gbb.12894

8. Nielsen CS, Staud R, Price DD. Individual Differences in Pain Sensitivity: Measurement, Causation, and Consequences. J Pain. 2009;10: 231–237. doi:10.1016/j.jpain.2008.09.010

9. Hestehave S, Abelson KSP, Brønnum Pedersen T, Munro G. The analgesic efficacy of morphine varies with rat strain and experimental pain model: implications for target validation efforts in pain drug discovery. Eur J Pain. 2019;23: 539–554. doi:10.1002/ejp.1327

10. Liang D-Y, Liao G, Wang J, Usuka J, Guo Y, Peltz G, et al. A Genetic Analysis of Opioid-induced Hyperalgesia in Mice. Anesthesiology. 2006;104: 1054–1062. doi:10.1097/00000542-200605000-00023

11. Duffy EP, Ward JO, Hale LH, Brown KT, Kwilasz AJ, Mehrhoff EA, et al. Sex and genetic background influence intravenous oxycodone self-administration in the hybrid rat diversity panel. Front Psychiatry. 2024;15: 1505898. doi:10.3389/fpsyt.2024.1505898

12. Mas M, Sabater E, Olaso MJ, Horga JF, Faura CC. Genetic variability in morphine sensitivity and tolerance between different strains of rats. Brain Res. 2000;866: 109–115. doi:10.1016/S0006-8993(00)02255-1

13. LaCroix-Fralish ML, Mogil JS. Progress in Genetic Studies of Pain and Analgesia. Annu Rev Pharmacol Toxicol. 2009;49: 97–121. doi:10.1146/annurev-pharmtox-061008-103222

14. Lüscher C, Robbins TW, Everitt BJ. The transition to compulsion in addiction. Nat Rev Neurosci. 2020;21: 247–263. doi:10.1038/s41583-020-0289-z

15. Gaddis N, Mathur R, Marks J, Zhou L, Quach B, Waldrop A, et al. Multi-trait genome-wide association study of opioid addiction: OPRM1 and beyond. Sci Rep. 2022;12: 16873. doi:10.1038/s41598-022-21003-y

16. Polimanti R, Walters RK, Johnson EC, McClintick JN, Adkins AE, Adkins DE, et al. Leveraging genome-wide data to investigate differences between opioid use vs. opioid dependence in 41,176 individuals from the Psychiatric Genomics Consortium. Mol Psychiatry. 2020;25: 1673–1687. doi:10.1038/s41380-020-0677-9

17. Zhou H, Rentsch CT, Cheng Z, Kember RL, Nunez YZ, Sherva RM, et al. Association of *OPRM1* Functional Coding Variant With Opioid Use Disorder: A Genome-Wide Association Study. JAMA Psychiatry. 2020;77: 1072. doi:10.1001/jamapsychiatry.2020.1206

18. Yang Y, Guan B, Wei Q, Wang W, Meng A. Morphine analgesia in male inbred genetic diversity mice recapitulates the among-individual variance in response to morphine in humans. Anim Models Exp Med. 2022;5: 288–296. doi:10.1002/ame2.12234

19. Belknap JK, Lame M, Danielson PW. Inbred strain differences in morphine-induced analgesia with the hot plate assay: A reassessment. Behav Genet. 1990;20: 333–338. doi:10.1007/BF01067800

20. Kest B, Hopkins E, Palmese CA, Adler M, Mogil JS. Genetic variation in morphine analgesic tolerance. Pharmacol Biochem Behav. 2002;73: 821–828. doi:10.1016/S0091-3057(02)00908-5

21. Nguyen JD, Grant Y, Taffe MA. Paradoxical changes in brain reward status during oxycodone self-administration in a novel test of the negative reinforcement hypothesis. Br J Pharmacol. 2021;178: 3797–3812. doi:10.1111/bph.15520

22. Kallupi M, De Guglielmo G, Carrette LLG, Simpson S, Kononoff J, Kimbrough A, et al. Large-scale behavioral characterization of oxycodone self-administration in heterogeneous stock rats reveals initial analgesic effects are associated with addiction-like behaviors. Neuropsychopharmacology. 2026 [cited 11 Feb 2026]. doi:10.1038/s41386-026-02348-8

23. Blackwood CA, McCoy MT, Ladenheim B, Cadet JL. Escalated Oxycodone Self-Administration and Punishment: Differential Expression of Opioid Receptors and Immediate Early Genes in the Rat Dorsal Striatum and Prefrontal Cortex. Front Neurosci. 2020;13: 1392. doi:10.3389/fnins.2019.01392

24. Sharp BM, Leng S, Huang J, Jones C, Williams RW, Chen H. Inbred rat heredity and sex affect oral oxycodone self-administration and augmented intake in long sessions: correlations with anxiety and novelty-seeking. Kavushansky A, editor. PLOS ONE. 2025;20: e0314777. doi:10.1371/journal.pone.0314777

25. Kuhn BN, Cannella N, Crow AD, Lunerti V, Gupta A, Walterhouse SJ, et al. Distinct Behavioral Profiles and Neuronal Correlates of Heroin Vulnerability Versus Resiliency in a Multi-Symptomatic Model of Heroin Use Disorder in Rats. Am J Psychiatry. 2025;182: 198–208. doi:10.1176/appi.ajp.20230623

26. Leonardo M, Brunty S, Huffman J, Kastigar A, Dickson PE. Intravenous fentanyl self-administration in male and female C57BL/6J and DBA/2J mice. Sci Rep. 2023;13: 799. doi:10.1038/s41598-023-27992-8

27. Beierle JA, Yao EJ, Goldstein SI, Scotellaro JL, Sena KD, Linnertz CA, et al. Genetic basis of thermal nociceptive sensitivity and brain weight in a BALB/c reduced complexity cross. Mol Pain. 2022;18: 17448069221079540. doi:10.1177/17448069221079540

28. Mogil JS, Wilson SG. Nociceptive and morphine antinociceptive sensitivity of 129 and C57BL/6 inbred mouse strains: Implications for transgenic knock-out studies. Eur J Pain. 1997;1: 293–297. doi:10.1016/S1090-3801(97)90038-0

29. Liu SX, Gades MS, Swain Y, Ramakrishnan A, Harris AC, Tran PV, et al. Repeated morphine exposure activates synaptogenesis and other neuroplasticity-related gene networks in the dorsomedial prefrontal cortex of male and female rats. Drug Alcohol Depend. 2021;221: 108598. doi:10.1016/j.drugalcdep.2021.108598

30. Wala EP, Holtman JR. Buprenorphine-induced hyperalgesia in the rat. Eur J Pharmacol. 2011;651: 89–95. doi:10.1016/j.ejphar.2010.10.083

31. Wang H, Akbar M, Weinsheimer N, Gantz S, Schiltenwolf M. Longitudinal Observation of Changes in Pain Sensitivity during Opioid Tapering in Patients with Chronic Low-Back Pain. Pain Med. 2011;12: 1720–1726. doi:10.1111/j.1526-4637.2011.01276.x

32. Li X, Angst MS, Clark JD. A murine model of opioid-induced hyperalgesia. Mol Brain Res. 2001;86: 56–62. doi:10.1016/S0169-328X(00)00260-6

33. Elhabazi K, Ayachi S, Ilien B, Simonin F. Assessment of Morphine-induced Hyperalgesia and Analgesic Tolerance in Mice Using Thermal and Mechanical Nociceptive Modalities. J Vis Exp. 2014; 51264. doi:10.3791/51264

34. Larson CM, Barajas C, Kitto KF, Wilcox GL, Fairbanks CA, Peterson CD. Development of opioid analgesic tolerance in rat to extended-release buprenorphine formulated for laboratory subjects. Tang S-J, editor. PLOS ONE. 2024;19: e0298819. doi:10.1371/journal.pone.0298819

35. Laboureyras E, Boujema MB, Mauborgne A, Simmers J, Pohl M, Simonnet G. Fentanyl-induced hyperalgesia and analgesic tolerance in male rats: common underlying mechanisms and prevention by a polyamine deficient diet. Neuropsychopharmacology. 2022;47: 599– 608. doi:10.1038/s41386-021-01200-5

36. Larson CM, Peterson CD, Kitto KF, Wilcox GL, Fairbanks CA. Sustained-release buprenorphine induces acute opioid tolerance in the mouse. Eur J Pharmacol. 2020;885: 173330. doi:10.1016/j.ejphar.2020.173330

37. Bryant CD, Roberts KW, Byun JS, Fanselow MS, Evans CJ. Morphine analgesic tolerance in 129P3/J and 129S6/SvEv mice. Pharmacol Biochem Behav. 2006;85: 769–779. doi:10.1016/j.pbb.2006.11.012

38. Pantouli F, Grim TW, Schmid CL, Acevedo-Canabal A, Kennedy NM, Cameron MD, et al. Comparison of morphine, oxycodone and the biased MOR agonist SR-17018 for tolerance and efficacy in mouse models of pain. Neuropharmacology. 2021;185: 108439. doi:10.1016/j.neuropharm.2020.108439

39. Chia Y-Y, Liu K, Wang J-J, Kuo M-C, Ho S-T. Intraoperative high dose fentanyl induces postoperative fentanyl tolerance. Can J Anesth Can Anesth. 1999;46: 872–877. doi:10.1007/BF03012978

40. Guignard B, Bossard AE, Coste C, Sessler DI, Lebrault C, Alfonsi P, et al. Acute Opioid Tolerance: Intraoperative Remifentanil Increases Postoperative Pain and Morphine Requirement. Anesthesiology. 2000;93: 409–417. doi:10.1097/00000542-200008000-00019

41. Chu LF, Clark DJ, Angst MS. Opioid Tolerance and Hyperalgesia in Chronic Pain Patients After One Month of Oral Morphine Therapy: A Preliminary Prospective Study. J Pain. 2006;7: 43–48. doi:10.1016/j.jpain.2005.08.001

42. Hoffmann O, Plesan A, Wiesenfeld-Hallin Z. Genetic differences in morphine sensitivity, tolerance and withdrawal in rats. Brain Res. 1998;806: 232–237. doi:10.1016/S0006-8993(98)00768-9

43. Madia PA, Dighe SV, Sirohi S, Walker EA, Yoburn BC. Dosing protocol and analgesic efficacy determine opioid tolerance in the mouse. Psychopharmacology (Berl). 2009;207: 413–422. doi:10.1007/s00213-009-1673-6

44. Mogil JS, Wilson SG, Bon K, Eun Lee S, Chung K, Raber P, et al. Heritability of nociception I: Responses of 11 inbred mouse strains on 12 measures of nociception. Pain. 1999;80: 67–82. doi:10.1016/S0304-3959(98)00197-3

45. Lariviere WR, Wilson SG, Laughlin TM, Kokayeff A, West EE, Adhikari SM, et al. Heritability of nociception. III. Genetic relationships among commonly used assays of nociception and hypersensitivity. Pain. 2002;97: 75–86. doi:10.1016/S0304-3959(01)00492-4

46. Nielsen CS, Stubhaug A, Price DD, Vassend O, Czajkowski N, Harris JR. Individual differences in pain sensitivity: Genetic and environmental contributions. Pain. 2008;136: 21–29. doi:10.1016/j.pain.2007.06.008

47. Angst MS, Phillips NG, Drover DR, Tingle M, Ray A, Swan GE, et al. Pain sensitivity and opioid analgesia: A pharmacogenomic twin study. Pain. 2012;153: 1397–1409. doi:10.1016/j.pain.2012.02.022

48. Volkow ND, McLellan AT. Opioid Abuse in Chronic Pain — Misconceptions and Mitigation Strategies. Longo DL, editor. N Engl J Med. 2016;374: 1253–1263. doi:10.1056/NEJMra1507771

